# The tilt illusion arises from an efficient reallocation of neural coding resources at the contextual boundary

**DOI:** 10.1101/2024.09.17.613538

**Authors:** Ling-Qi Zhang, Jiang Mao, Geoffrey K. Aguirre, Alan A. Stocker

## Abstract

The tilt illusion — a bias in the perceived orientation of a center stimulus induced by an oriented surround — illustrates how context shapes visual perception. While the tilt illusion has been the subject of quantitative study for over 85 years, we still lack a comprehensive account of the phenomenon that connects its neural and behavioral characteristics. Here, we demonstrate that the tilt illusion originates from a dynamic change in neural coding precision induced by the surround context. We simultaneously obtained psychophysical and fMRI responses from human subjects while they viewed gratings in the absence and presence of an oriented surround, and extracted sensory encoding precision from their behavioral and neural data. Both measures show that in the absence of a surround, encoding reflects the natural scene statistics of orientation. However, in the presence of an oriented surround, encoding precision is significantly increased for stimuli similar to the surround orientation. This local change in encoding is sufficient to accurately predict the behavioral characteristics of the tilt illusion using a Bayesian observer model. The effect of surround modulation increases along the ventral stream, and is localized to the portion of the visual cortex with receptive fields at the center-surround boundary. The pattern of change in coding accuracy reflects the surround-conditioned orientation statistics in natural scenes, but cannot be explained by local stimulus configuration. Our results suggest that the tilt illusion naturally emerges from a dynamic coding strategy that efficiently reallocates neural coding resources based on the current stimulus context.

## Introduction

Human perception is significantly influenced by sensory context. A classic demonstration is the tilt illusion, in which the perceived orientation of a center stimulus is altered by the orientation of a surround^1^. Previous investigations of the tilt illusion have mainly focused on how surround context alters the response characteristics of individual neurons. For example, orientation-selective neurons in early visual cortex both suppress their responses close to, and shift their tuning preferences away from, the contextual orientation^2,3,4,5^. The non-classical receptive field (RF) is a closely related phenomenon in which neural responses evoked by stimuli within the RF exhibit complex dependencies upon content outside the RF^6,7^. These surround-dependent changes in neural response have been attributed to divisive gain control^8^, which removes redundancies in neural signals by a normalization mechanism^4,9^.

Connecting these observations at the level of single neurons to perceptual behavior, however, is challenging. Doing so requires specific assumptions regarding how sensory information is both represented (i.e., encoded) and interpreted (i.e., decoded) by neural populations at different processing stages across the sensorimotor stream. Practically, it also requires the difficult task of recording from large populations of sensory neurons under contextual modulation. Therefore, previous modeling work has approached this problem by relying on simulated neural population responses instead^10,11^. Generally, we lack a coherent theoretical framework that provides a functional and teleological account of the tilt illusion at the level of the observer, and quantitatively connects empirical measures of neural population responses and behavior.

Here, we provide this synthesis by studying orientation perception with simultaneous measurements of psychophysical behavior and neural activity using functional Magnetic Resonance Imaging (fMRI). We analyzed these data within an information-theoretic framework. Specifically, we extracted the Fisher information (FI) of orientation encoding as a measure of encoding accuracy from both behavioral responses and neural activity patterns. We computed “behavioral FI” based on a lawful relationship between FI and the bias and variance of psychophysical stimulus estimates^12,13^. We also obtained “neural FI” for early visual areas by fitting voxel-wise probabilistic encoding models to the fMRI data^14,15^. Within this framework, behavioral and neural measures of encoding accuracy can be directly compared to each other, and (via the efficient coding hypothesis) to orientation priors measured from natural scenes^16,17^. Furthermore, we can leverage the retinotopic organization of occipital cortex to determine where potential changes in neural encoding accuracy arise relative to the spatial structure of the stimulus.

Our results show that neural and behavioral measures of encoding accuracy are qualitatively similar across all conditions tested. In the absence of an oriented surround, orientation encoding precision reflects the orientation statistics of natural scenes. However, in the presence of a spatially oriented surround, encoding accuracy significantly increases at the surround orientation in a way consistent with the conditional orientation statistics of spatially adjacent regions of natural scenes. The changes in encoding, however, cannot be explained by local effects of stimulus configuration (i.e., “vignetting”). We further demonstrate that the change in encoding precision measured at the neural level is sufficient to fully predict observer perceptual reports of the tilt illusion based on a Bayesian observer model of orientation estimation^18^. Finally, we find that the change in neural encoding occurs at the boundary between the center and surround of the stimulus, with its magnitude increasing along the ventral visual hierarchy. Our results support the notion that the tilt illusion arises from an efficient reallocation of coding resources based on stimulus context.

## Results

We conducted a delayed orientation estimation task inside the fMRI scanner while measuring blood-oxygen-level-dependent (BOLD) activity (Fig. 1A). Trials began with a 1.5 second presentation of a grating stimulus. The grating was presented within an annular surround consisting of either non-oriented noise (baseline), or a grating with one of two fixed orientations (± 35 degrees off vertical). Following a brief, blank delay, a probe stimulus (line) appeared, and subjects were asked to rotate the probe using a two-button response pad to report their perceived orientation of the grating. Every block of the fMRI acquisition consisted of 20 trials in one of the three fixed surround conditions. The order of surround conditions was randomized and counterbalanced across acquisitions. Each subject completed a total of 1,200 trials across six sessions of data collection.

**Figure 1:**
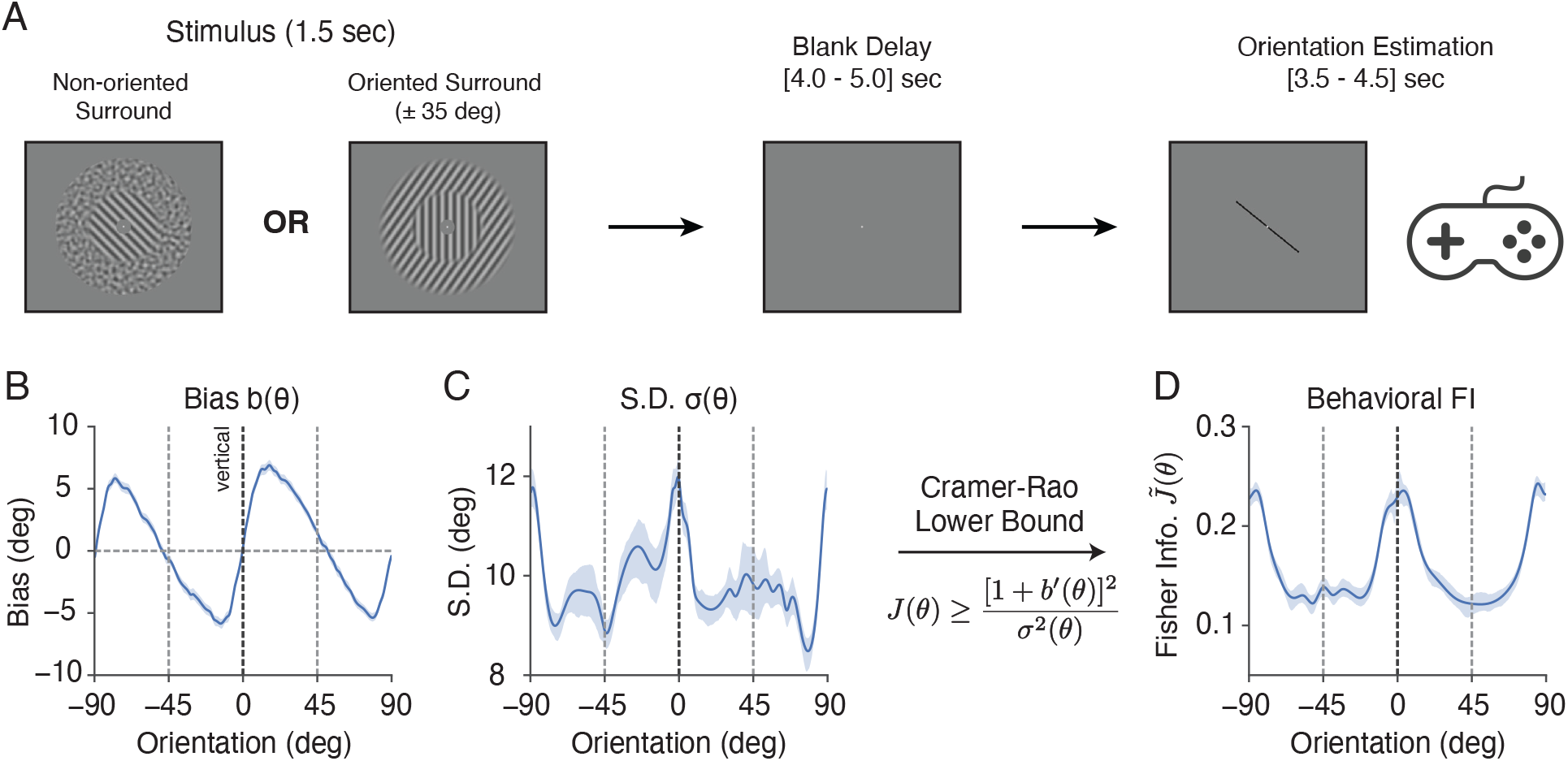
Experimental design and behavioral data analysis. **A)** Subjects (n = 10) performed a delayed orientation estimation task across 1,200 trials during fMRI. Target (center) gratings were presented within an annular surround of either non-oriented, spatially filtered noise pattern, or one of two fixed grating orientations (±35 degrees off vertical). **B) - D)** Behavioral data for the combined subject in the non-oriented surround condition; see Supplementary Fig. 6 for individual subjects. **B)** Estimation bias *b*(*θ*) as a function of stimulus orientation. **C)** Standard deviation *σ*(*θ*) of the estimates as a function of stimulus orientation. **D)** Fisher Information (square root, normalized; denoted as 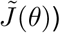 quantifying orientation encoding precision, derived from estimation bias and variance using the Cramer-Rao Lower Bound^19,13^ (see *Methods*). Shaded areas indicate ±SEM.

### Behavioral measure of orientation encoding accuracy

We first examined the perceptual behavior of subjects in the non-oriented surround (baseline) condition. Fig. 1B depicts the estimation bias *b*(*θ*) as a function of the target orientation. Estimates exhibited a well-known oblique bias, i.e., the perceived orientation of the grating was biased away from cardinal (i.e., vertical and horizontal) orientations^20,21,22,13^. For example, when the target was slightly rotated clockwise (positive) from the vertical, the bias was positive, indicating that subjects perceived the orientation to be even more clockwise. Additionally, the standard deviation (SD) of the estimates *σ*(*θ*) was higher at cardinal compared to oblique orientations (Fig. 1C).

We took advantage of the Cramer–Rao Lower Bound (CRLB) to quantify encoding accuracy based on behavioral data^19^. The CRLB describes the bounded, lawful relationship between estimation bias *b*(*θ*) and variance *σ*^2^(*θ*) of an estimator, and the FI *J*(*θ*) of its sensory encoding as follows (also see *Methods*):

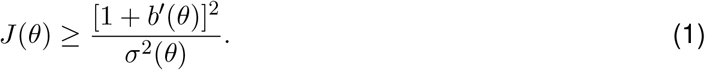

We have previously demonstrated^13^ that equating this lower bound with FI requires only the weak and common assumption that the estimator and the subsequent response process (i.e., motor control) are not corrupted by stimulus-dependent noise. Therefore, Eq.1 allows us to extract encoding accuracy in terms of FI from subject responses without the need to assume a specific decoding model.

We extracted this behavioral FI from the bias and variance data. We found that orientation encoding in the non-oriented surround condition was non-homogeneous (Fig 1D): FI was highest at the cardinal orientations, and lowest at the obliques. Because FI is inversely related to discriminability^11,23,12^, our result is consistent with previous measurements of orientation discrimination thresholds, which have consistently shown lower thresholds at cardinal than oblique orientations^21,24^.

### Neural measure of orientation encoding accuracy

Next, we extracted neural measures of encoding accuracy from BOLD fMRI signals recorded during the delay period. We defined regions of interest (ROIs) based on separately measured retinotopic maps for each subject. Voxels from different visual areas within the visual eccentricity range of the grating stimulus were selected. We first fit a voxel-wise probabilistic encoding model^14,15^ to the normalized BOLD activity, averaged between 4 and 8 seconds for each trial after stimulus onset. Separate models were fit for each ROI, each subject, and each surround condition. The encoding model describes the activity of each voxel as a weighted sum of responses from a set of basis functions. Additionally, the model incorporates two sources of Gaussian noise: tuning-dependent noise and voxel-wise residual noise. Collectively, this model defines a multivariate voxel population encoding model *p*(**m**|*θ*) (Fig. 2A; see *Methods*, Eq. 11).

**Figure 2:**
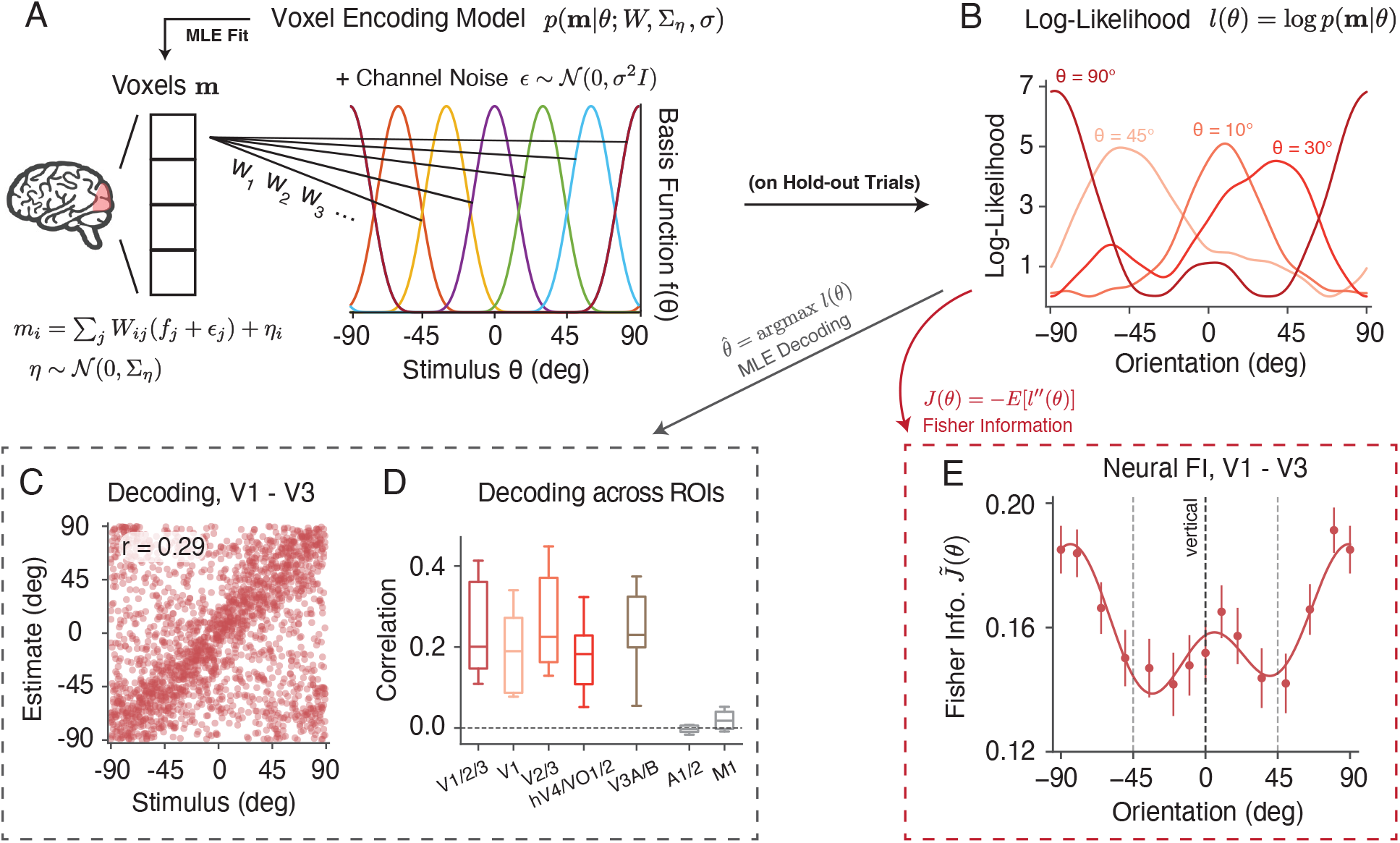
Neural data analysis. **A)** We described the voxel responses **m** using a population encoding model^14,15^, denoted as *p*(**m**|*θ*). The normalized activity for each voxel *m*_*i*_ was modeled as a weighted sum of responses from a set of basis functions. We assumed that each basis function was corrupted by channel noise *ϵ*. Additional variability of each voxel was modeled by residual noise *η*. Model parameters were obtained by fitting the voxel data using a two-stage procedure. **B)** The orientation log-likelihood of the model *l*(*θ*) = log *p*(**m**|*θ*) was obtained based on hold-out trials. **C)** We could decode the orientation of the stimulus presented on each trial as the orientation with highest likelihood, thus 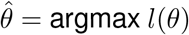. Here we show a scatter plot of the stimuli orientation (x-axis) versus the decoded orientation (y-axis) from the early visual cortex (V1 to V3), for all trials in the non-oriented surround condition from five subjects. **D)** Decoding correlation from different ROIs in the visual cortex and two control ROIs (auditory cortex and primary motor cortex). The box extends from the first to the third quartile of the average decoding correlation of all trials across individual subjects, with the center line at the median. The whiskers indicate the farthest data point. **E)** Fisher information (FI) of neural encoding was defined as the negative average second derivative of the log-likelihood, *J*(*θ*) = −*E*[*l*^′′^(*θ*)]. Shown is the neural FI (normalized, square root) of early visual cortex (V1 - V3) for the combined subject in the non-oriented surround condition. Error bars indicate ±SEM. See *Methods* for details.

For any given pattern of voxel BOLD activity **m**, the encoding model defined a sensory log-likelihood function *l*(*θ*) = log *p*(**m**|*θ*) (Fig. 2B). Previous studies have used the likelihood function to decode both the stimulus and its associated uncertainty for orientation^14^, motion direction^25^, and working memory content^26^ from BOLD activity. While not our main focus, we found that we could reliably decode orientation from a range of visual areas, but not from two control (auditory and motor) areas (Fig. 2C,D). Decoding performance was comparable between the three surround conditions, but tended to be slightly higher for the oriented surround (Supplementary Fig. S1).

We used the orientation log-likelihood to derive neural FI. For each trial, we calculated the negative second derivative of the log-likelihood function at the stimulus orientation (Fig. 2B). The neural FI is the expected value of this negative second derivative. To compute the FI for the combined subjects, we aggregated the results from individual participants and calculated the average within a 25-degree window centered at various orientations (Fig. 2E; see *Methods* for details). Consistent with behavioral FI (Fig. 1D), neural FI in the early visual cortex (V1 - V3) was highest around the cardinal and lowest close to the oblique orientations, in the non-oriented surround condition.

### Efficient encoding of orientation

In the previous sections, we demonstrated that in the non-oriented surround (baseline) condition both behavioral and neural measures of FI show similar, non-uniform patterns as a function of orientation. What is the origin of this non-homogenoues encoding pattern, and in particular, the emphasis for cardinal orientations? The efficient coding hypothesis suggests that there is a direct relationship between stimulus prior *p*(*θ*) and encoding FI for neural codes that aim for an optimal stimulus representation given resource constraints^16,17,27^:

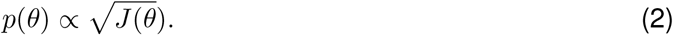

Thus, the normalized, square root of FI,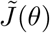, can be interpreted as the inferred orientation prior assuming an efficient neural encoding^29^, which allows for a direct comparison with other estimates of the orientation prior. For example, Figure 3A shows the statistics of local visual orientation computed over large subsets of photographic images containing more or fewer natural objects^28^. In both natural and human-made environments, the prior probability of cardinal orientations is higher than that of oblique orientations.

**Figure 3:**
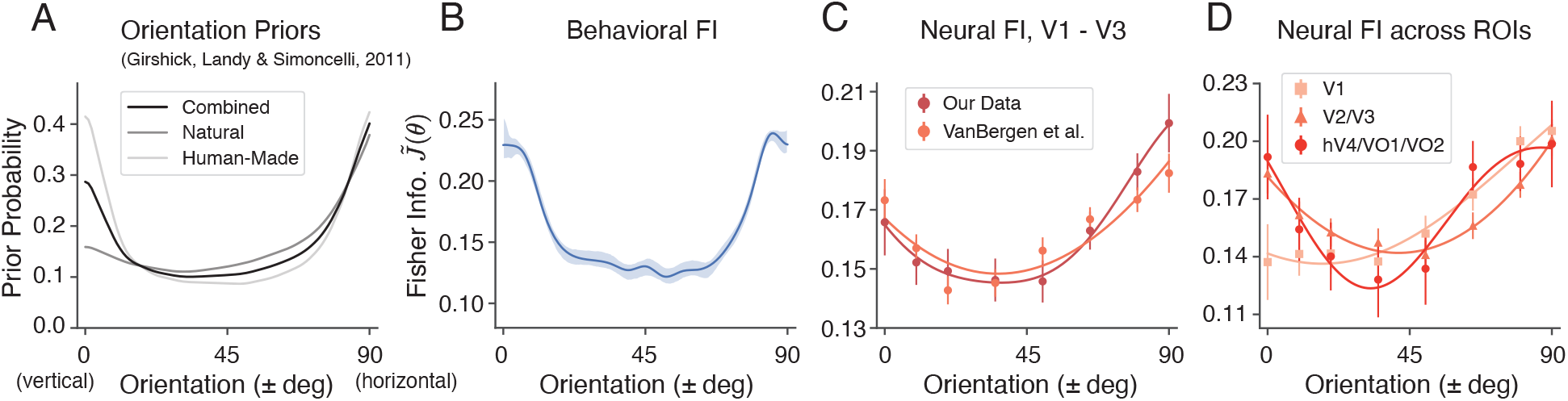
Comparison between the orientation priors derived from photographic images, the behavioral FI, and the neural FI in the non-oriented surround condition. For all panels, we assumed vertical symmetry and combined the data from corresponding counter-clockwise and clockwise orientations. **A)** Orientation priors measured in different visual environments, reproduced from Girshick et al.^28^. **B)** Behavioral FI calculated from the estimation data (same as in Fig. 1D). **C)** Neural FI in the early visual cortex calculated from the voxel encoding model for our data (Fig. 2E), and another dataset^14^. **D)** Neural FI for different visual areas. The data plotted are for the combined subject, shaded area and error bars indicate ±SEM. All FI curves represent the normalized, square root of Fisher information,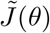.

We found that the behavioral FI pattern in the non-oriented surround condition resembled these environmental priors (Fig. 3B), which is consistent with the efficient coding hypothesis. Similarly, we observed the same qualitative match for the neural encoding accuracy (neural FI) in early visual cortex (V1 - V3), which was confirmed by the same analysis of a previously reported dataset^14^ (Fig. 3C). Lastly, to assess whether the orientation prior was reflected across different visual areas, we obtained the neural FI separately for three groups of ROIs, organized along the visual ventral hierarchy (Fig. 3D). We found a strong cardinal emphasis in the neural FI of areas V2 and V3, and hV4 and VO1/2; the neural FI in these areas was most similar to the orientation prior in natural scenes.

### Surround modulation of orientation encoding

We now consider the tilt illusion by examining the behavioral and neural data from the oriented surround conditions (Fig. 4). As in the previous analysis, we assumed symmetry around the vertical meridian and also aggregated the data measured from the two symmetric surround orientation conditions (i.e., positive angles for the +35 deg condition and negative angles for the -35 deg condition). We denote the 90-degree orientation range (vertical to horizontal) containing the surround orientation as “near-surround”, and the opposite range as “far-surround”, respectively.

**Figure 4:**
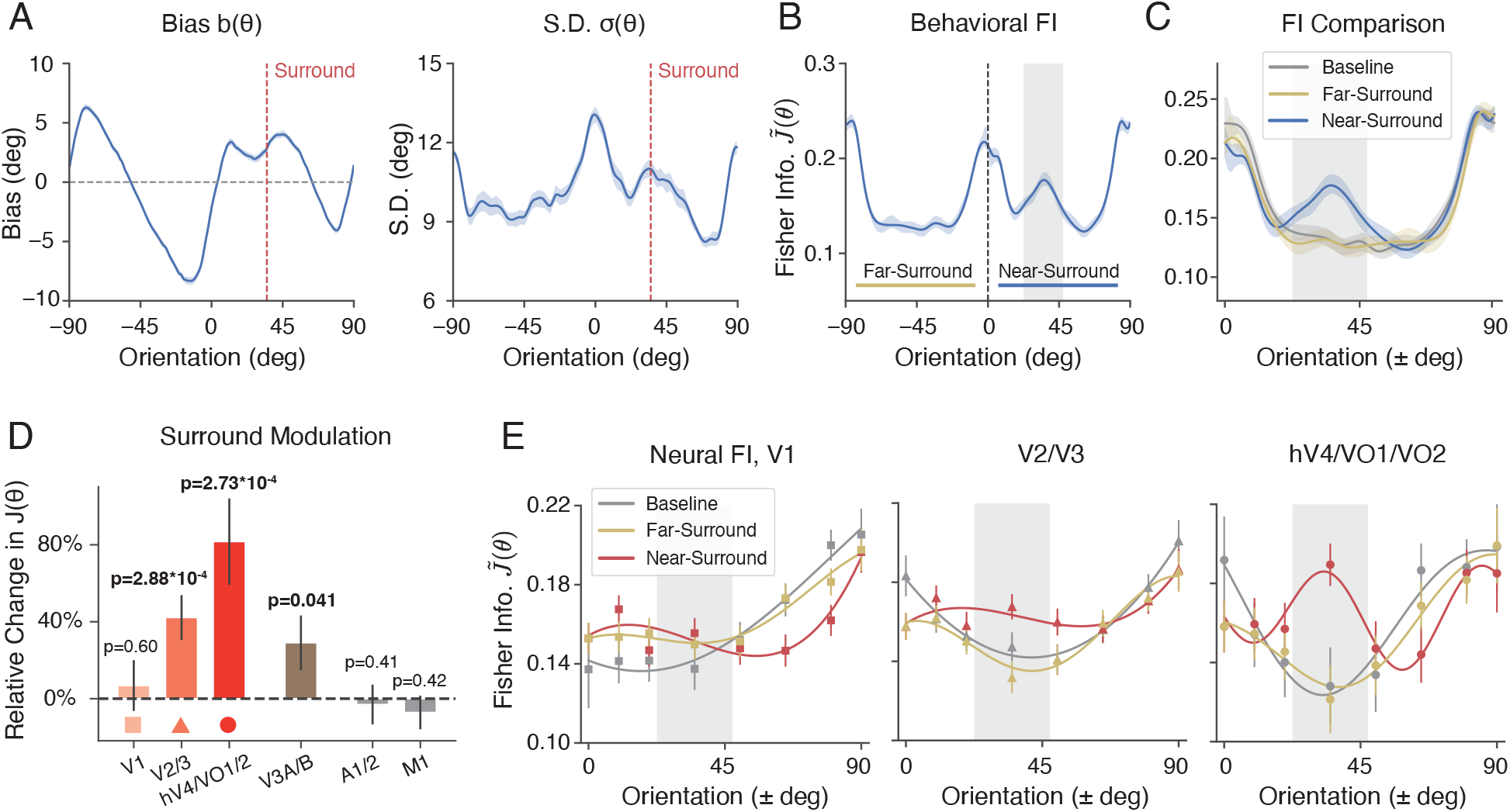
Orientation encoding in the tilt illusion. We analyzed the behavioral and neural data in the oriented surround condition in the same way as in the non-orientated condition before (combined subject). **A)** Estimation bias *b*(*θ*) and standard deviation *σ*(*θ*) as a function of the orientation of the center grating. The dashed line indicates the orientation of the surround. **B)** Behavioral FI, calculated from estimation data. “Near-surround” refers to the 90-degree orientation range (vertical to horizontal) on the side of the surround orientation, and “far-surround” refers to the 90-degree range on the side opposite to the surround orientation. Gray-shaded area indicates a 25-degree window (between 22.5 - 47.5 degree) centered at the surround orientation. **C)** Comparison of the behavioral FI between near-surround and far-surround orientations in the oriented and non-oriented (baseline) surround conditions. **D)** The relative percentage change in neural FI within the gray-shaded area, for different ROIs in the visual cortex, and two control ROIs. **E)** Comparison of neural FI along the visual ventral stream, between the near-surround side, far-surround side, and the baseline condition. Shaded areas and error bars indicate ±SEM. See Supplementary Fig. S6 and Supplementary Fig. S7 for comparison of behavioral and neural FI of individual subjects.

The oriented surround altered both the bias and variance of the orientation estimates of the center grating, especially for orientations close to the surround (compare Figs. 1B,C and Fig. 4A). The changes are consistent with well-known characteristics of the tilt-illusion^1,30^, showing a strong repulsive bias near the surround orientation and a subtle attractive bias further away (see also Supplementary Fig. S2A). We again used Eq.1 to extract behavioral FI from the estimation bias and standard deviation shown in Fig. 4A. We found that the oriented surround leads to a significant increase in encoding precision close to the surround orientation, while the overall FI pattern – in particular for the “far surround” range – remains unchanged (Fig. 4B). This is particularly apparent when plotting the behavioral FI for both the near- and far-surround range alongside the FI for the non-oriented surround (baseline) condition (Fig. 4C).

This characteristic change in orientation encoding is also present at the neural level. We derived the neural FI by fitting a separate set of voxel encoding models to the fMRI data collected in the oriented surround condition. We found a significant effect of surround modulation on encoding accuracy in several areas of early visual cortex. Consistent with the behavioral measure, neural FI is substantially increased within a narrow window near the surround orientation as compared to the non-oriented surround (baseline) condition (see *Methods*). The magnitude of this effect increases along the visual ventral stream with an apparent peak in the combined area hV4/V01/V02 (Fig. 4D, Fig. S3A-B). Particularly in these latter areas, the encoding pattern (Fig. 4E) are remarkably similar to the behavioral FI (Fig. 4C).

We further examined how the change in neural encoding precision depended upon which part of the stimulus was encoded. We computed neural FI for different subsets of voxels with different eccentricity ROIs based on the center and size of their population receptive fields (pRF; see *Methods*). We found that encoding precision computed for voxels with pRFs exclusively within the surround region did not exhibit any effect of surround modulation (Fig. 5C, > 9 and > 15). Similarly, encoding precision extracted for voxels with pRFs strictly within the center remained unaffected by the surround (Fig. 5C, < 5 and < 1.5). In contrast, encoding precision for voxels at the contextual boundary were strongly modulated (Fig. 5C, 5 - 9). This suggests that changes in neural FI are driven by modulation of the center encoding through interactions between center and surround regions, and that these modulations are spatially localized to the area close to the center-surround contextual boundary.

**Figure 5:**
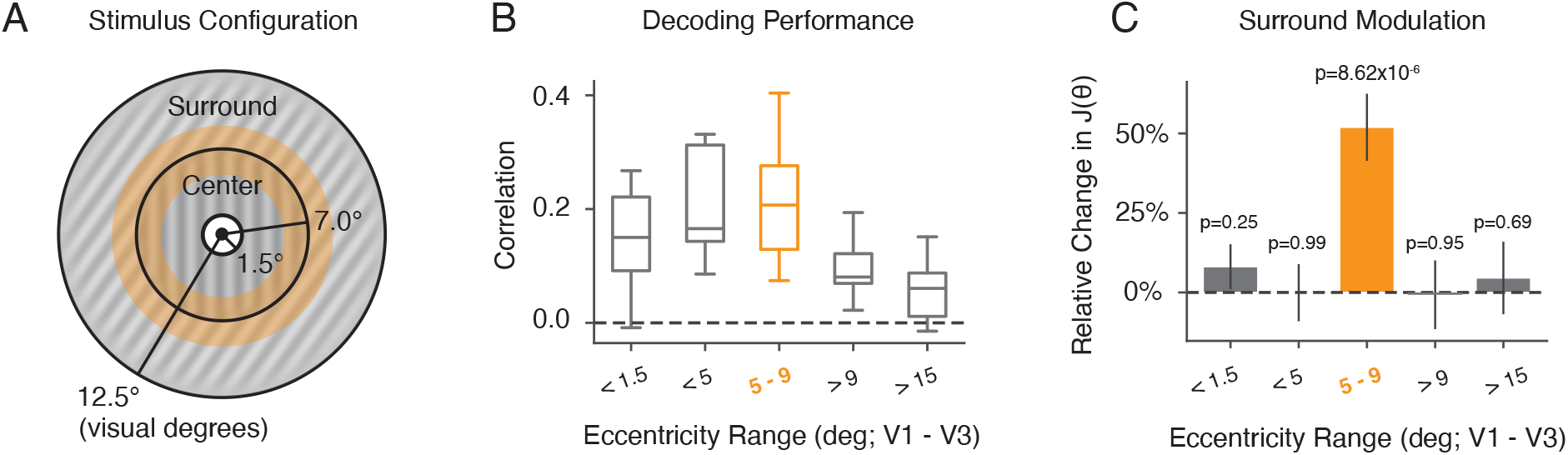
Surround modulation for ROIs at different stimulus eccentricities. **A)** Spatial configuration of center-surround stimuli used in our experiment. The center (target) extends from 1.5 to 7 degrees of visual angle in radius. The surround extends from 7 to 12.5 degrees radius. The orange area marks the center-surround contextual boundary (5 to 9 degrees). **B) - C)** Voxels from within area V1 - V3 were selected based on the center and size of their pRFs (see *Methods*). **B)** Average decoding correlation for all subjects using voxels with ROIs at different stimulus eccentricities. **C)** The relative change in neural FI with respect to the baseline near the surround orientation for different ROI eccentricities. Error bars indicate ±SEM.

### Predicting the tilt illusion from neural encoding accuracy

So far, we have established a tight correspondence between behaviorally and neurally estimated encoding accuracy. We have shown that the tilt illusion coincides with a consistent, characteristic increase in encoding precision for orientations similar to the surround orientation. To demonstrate a causal role of these encoding changes, we tested whether the observed neural changes in FI can directly predict the psychophysical reports of the tilt illusion (Fig. 6A). We employed a recently developed Bayesian observer model for orientation estimation^18^. The model assumes that encoding is efficient (Eq. 2), which jointly constrains the model’s likelihood function and prior distribution. Thus, for any given function of the encoding precision (e.g., measured as FI) the model is tightly constrained and able to make quantitative predictions of subjects’ orientation estimates.

**Figure 6:**
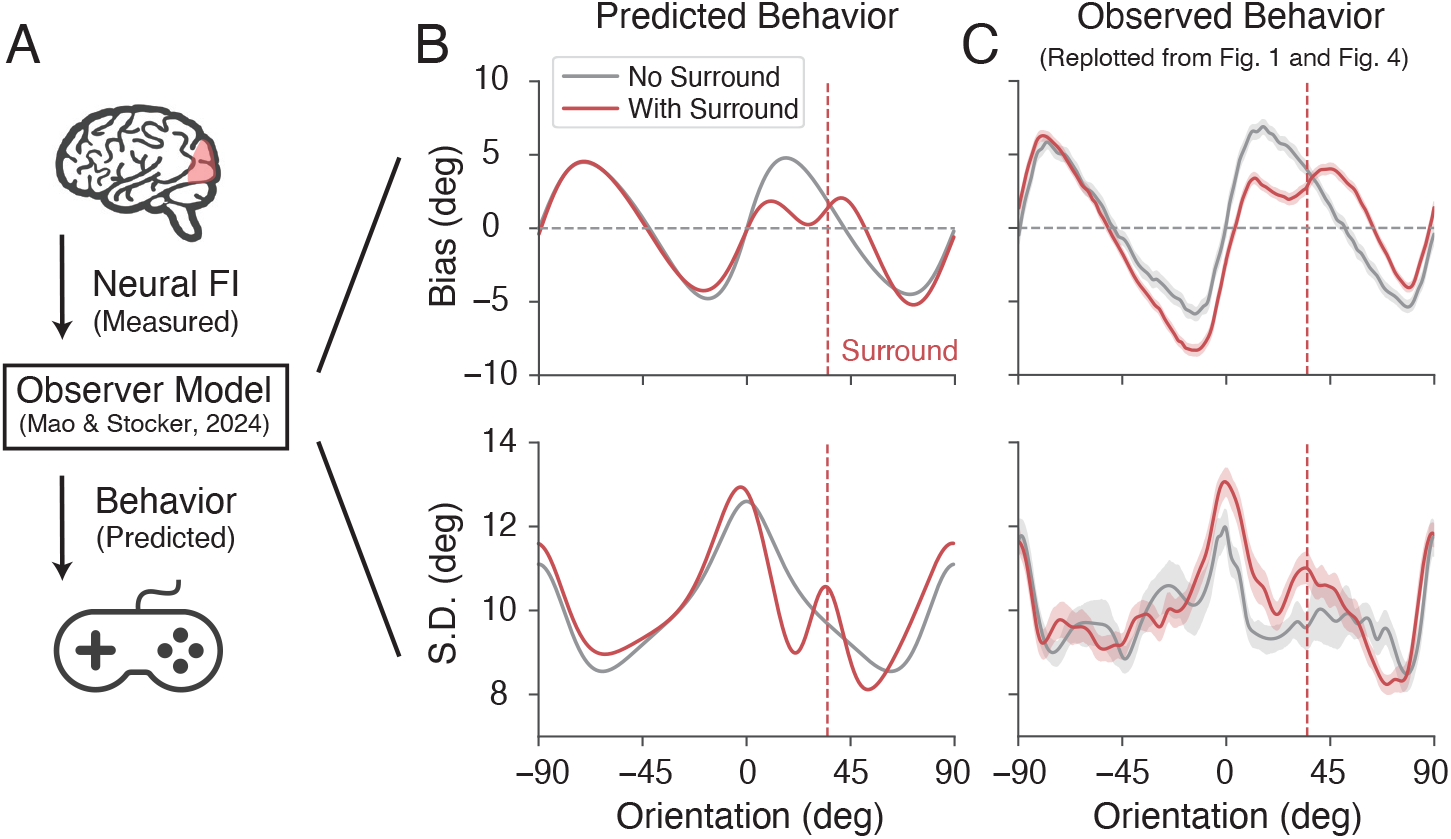
Predicting the tilt illusion from the neurally measured encoding precision. **A)** We used the extracted neural FI of areas hV4/VO1/VO2 in the non-oriented surround condition (baseline) and oriented surround condition to predict subjects’ behavior (i.e., mean and standard deviation of their orientation estimates) based on a recent state-of-the-art Bayesian observer model for orientation estimation^18^. Data and predictions are for the combined subject. **B)** Predicted bias and standard deviation. **C)** Measured estimation bias and standard deviation, replotted from Fig. 1 and Fig. 4. Gray curves indicate the baseline and red curves indicate the oriented surround condition. Shaded areas represent ±SEM.

We set the encoding precision of the model to reflect the neural FI measured for areas hV4/VO1/2 (Fig. 4). We first used the data from the baseline condition to determine the remaining free global parameters of the model (e.g., overall sensory noise). Then, we updated the modeled encoding precision to match the neural FI measured for the surround condition. The model output provided predictions of the perceptual bias and standard deviation in the absence and presence of an oriented surround (see *Methods* for more details.). As shown in Fig. 6B, the model successfully recapitulated the pattern of estimation bias and standard deviation in the baseline condition (gray lines), which confirms the result of the previous study^18^. Moreover, it accurately predicted the detailed, characteristic changes in bias and standard deviation observed in the tilt illusion (red lines). This included the repulsive bias near the surround orientation (as indicated by the positive slope of the bias curve; one of the most prominent features of the tilt illusion), as well as the accompanying increase in estimation SD (see also Supplementary Fig. S2).

Note that a key assumption of the model is that orientation reports are the result of a holistic inference process that jointly operates at low- and high-level representations of the stimulus (i.e., stimulus orientation, but also orientation categories, such as vertical and horizontal orientations). Here, we assumed that subjects also treat the surround orientation as an implicit category boundary. We verified that incorporating both the dynamic change in encoding precision and the categorical boundary at the surround are necessary for the model to make correct predictions of the tilt illusion effect (see Supplementary Fig. S4).

### Neural mechanism of surround modulation

We have demonstrated that the tilt illusion arises from changes in orientation encoding in the presence of an oriented surround context. What is the origin of these changes in encoding accuracy? One possibility is that the addition of an oriented surround naturally leads to increased coding accuracy near the surround orientation because of the nonlinear processing of the visual system. In this case, there are no changes in the response properties of sensory neurons, and the observed difference in encoding accuracy is purely due to the spatial configuration of the stimulus. Alternatively, the presence of a surround context actively alters the orientation response properties of sensory neurons^4,6^, resulting in the observed increase in coding precision.

The potential effect of spatial configuration is closely related to the issue of “stimulus vignetting”^32,33^, in which the arrangement of the stimulus and its aperture can result in additional signals for orientation decoding. To quantify the changes in the measured encoding FI that arise solely due to differences in stimulus configuration (i.e., random vs. oriented surround) in the absence of changes in neural responses properties, we implemented an image-computable voxel encoding model^33^. The model first applies a decomposition to the stimulus image, generating multiple bands of filter responses with varying orientations and spatial frequency selectivity (a steerable pyramid^34^).

A map of voxel responses can then be obtained by averaging across these bands. While each voxel in this construction is not orientation-selective, the pattern of responses across voxels as the grating rotates can still provide information about grating orientation. We again quantify this information using FI. (Fig. 7A, see *Methods*).

**Figure 7:**
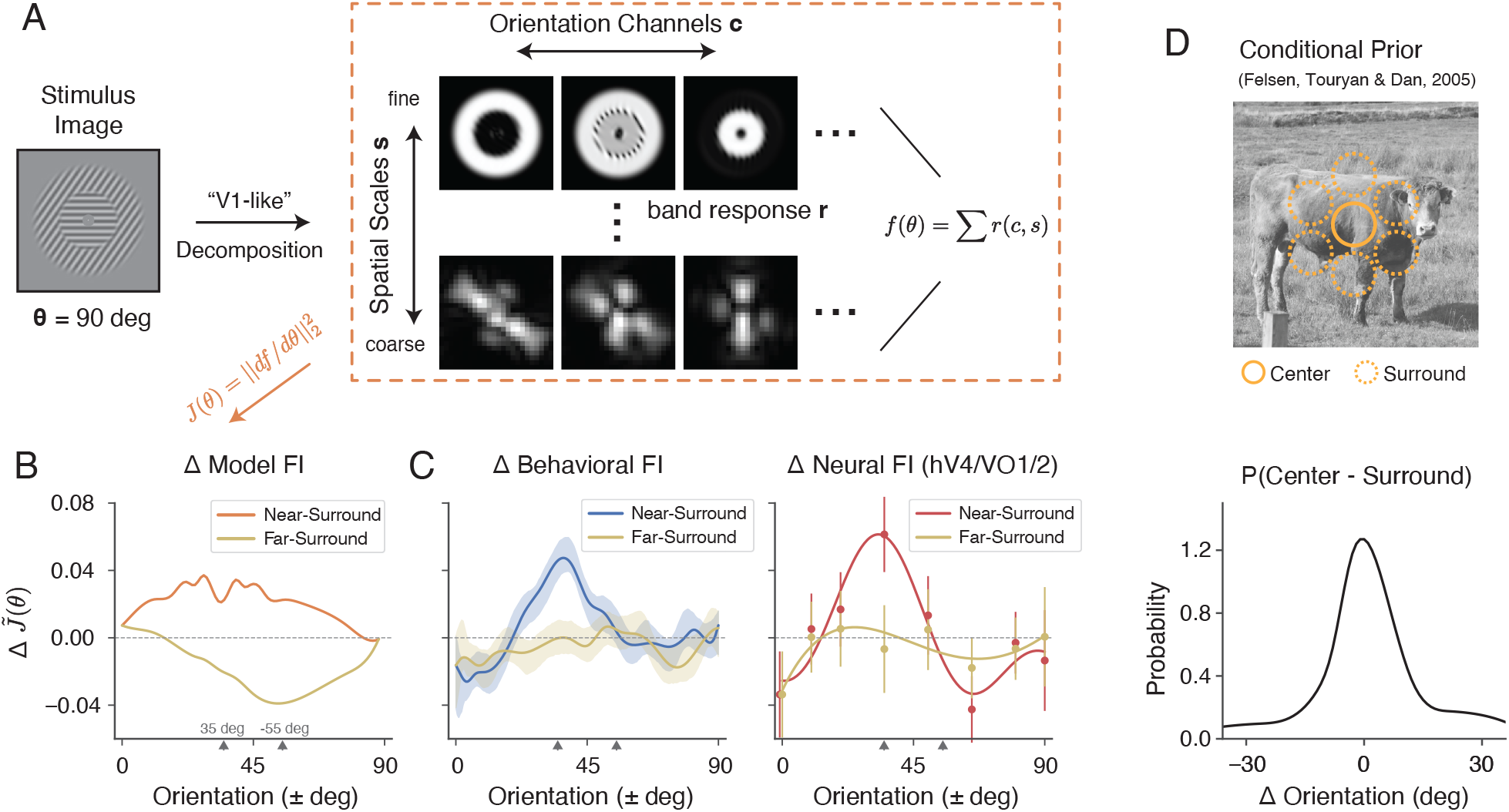
The effect of surround modulation cannot be explained by stimulus configuration, but is consistent with natural scene statistics. **A)** We simulated a “retinotopic map” of voxel responses *f* (*θ*) by averaging across different orientation channels and spatial scales in a steerable pyramid decomposition *r*(*c, s*) (see *Methods*). **B)** Changes in encoding FI 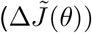) between stimuli with oriented surround compared to stimuli with random surround (baseline) condition based on the steerable pyramid voxel encoding model. The two ticks on the x-axis denote the surround orientation (+35 deg) and the orientation orthogonal to the surround (−55 deg). **C)** Changes in behavioral FI and neural FI with surround modulation compared to the baseline condition. **D)** Probability distribution of the angular difference in orientation between the center and surround regions of natural images (adapted from Felsen et al. ^31^). Shaded area and error bars indicate ± SEM.

For stimuli with a non-oriented surround, the encoding FI was non-zero (Supplementary Fig. S5), reflecting the vignetting effect reported by Roth et al.^33^. Note, however, that the FI is uniform across orientation because any effect of stimulus configuration in the non-oriented condition is isotropic by design. Next, we calculated the changes in FI for stimuli in the oriented surround condition (Fig. 7B). We found that the oriented surround elicited a broad increase in FI for the near-surround orientations compared to the baseline condition. At the same time, it also caused a broad decrease in FI for the far-surround orientations, with the lowest point at the orientation orthogonal to the surround (−55 degrees). This pattern was unlike the changes in FI we observed in both our behavioral and neural data (Fig. 7C): we observed an increase in FI that was limited to a small range around at the surround orientations, while encoding accuracy for far-surround orientations remained essentially unchanged. Thus, the effect of stimulus configuration cannot explain the measured changes in encoding accuracy. Rather, additional surround-induced mechanisms must be at work that dynamically adjust the neural representation of stimulus orientation, similar to what has been observed at the single-cell level^2,3,4,5,6^.

But why should the visual system actively increase encoding precision close to the surround orientation? Again, we turn to the efficient coding hypothesis, which suggests that the increase in FI should correspond to a local increase in the probability of those orientations. Spatial structures in adjacent regions of natural images are indeed correlated^35^. Therefore, the observation of a specific surround orientation indicates a marked increase in the probability of the center orientation being similar to that of the surround (Fig. 7D). We found that the change in encoding FI closely resembles the probability distribution of orientation difference between center and surround regions in natural images. The effective range of surround modulation is similar to the width of this distribution (compare Fig. 7C, D). We thus conclude that the effect of surround modulation is consistent with a form of dynamic efficient coding, in which coding resources are actively reallocated based on contextual information.

## Discussion

Our study reveals the sensory origin of the well-known tilt illusion. Based on concordant measures of encoding precision from behavioral and neural data, we demonstrate that the presence of an oriented surround causes a dynamic change in neural encoding precision, such that sensory representations remain optimized for both the long-term as well as the local surround-conditioned statistics of orientations found in natural scenes. The strength of the neural encoding change increases along the visual ventral stream, and is spatially localized to the boundary between the center and contextual surround. Furthermore, we show that the reported encoding change is sufficient to predict subjects’ behavior in the tilt illusion using a state-of-the-art Bayesian observer model of orientation estimation. Our findings support the notion that the tilt illusion is a manifestation of a sensory system that dynamically updates its encoding characteristics according to stimulus context in order to maximize information capacity^10,36,37,38,39^.

We use Fisher information as a common metric to quantify sensory encoding precision, which offers several advantages. First, it allows us to extract sensory encoding characteristics from participants’ reports in our psychophysical orientation estimation task using a lower-bound relation between FI and estimation bias and variance^13^. It also allows us to directly compare our results with discrimination threshold experiments, which directly quantify encoding precision, since discrimination thresholds are inversely proportional to FI^11^. Previous studies have reported discrimination thresholds with^40^ and without^21,24^ spatial context that are well aligned with our results. Finally, Fisher information enables a direct comparison of encoding accuracy derived from simultaneously recorded behavioral and neural data.

While the extracted Fisher information precisely quantifies how the precision of sensory encoding changes in the presence of an oriented surround, it does not specify the underlying neural mechanisms responsible for these changes^41^. Previous studies have documented a diverse set of possible mechanisms at the level of neuronal tuning characteristics including changes in response gain, tuning preference, and tuning width^5,7,31,42,43^. All these changes combined and accumulated across a neural population, as well as potential noise correlations^44,45^, then determine FI at the level that we have measured in our study. Thus, our results provide tight quantitative constraints for identifying the underlying neural mechanisms and their interactions across the population. Future research that involves recordings from large neural populations under contextual modulation will be necessary to more definitely establish connections between mechanisms operating at the individual neuron level and the population-wide changes in encoding precision we have found here.

Our results support converging lines of evidence suggesting that the sensory cortex forms efficient representations of perceptual variables according to their long-term (prior) statistics in natural scenes. For example, Harrison et al.^46^ used electroencephalogram (EEG) measurements and a forward encoding model to show that the tuning properties of cortical neurons can encode an orientation prior. Similarly, based on single-unit recording data, Zhang and Stocker^29^ illustrated that a power-law, slow speed prior for visual motion is represented in macaque MT cortex via a logarithmic encoding mechanism. What sets our results apart from these previous findings is that they are obtained from a joint analysis of simultaneously recorded behavioral and neural data. Whole brain fMRI recordings also allowed us to pinpoint and track the neural representation of orientation priors across the representational hierarchy of human visual cortex.

Furthermore, we show that the context-induced changes in neural encoding ensure that the sensory representation remains efficient with regard to the natural orientation statistics conditioned on the dominant surround orientation (Fig. 5). This offers a new normative understanding of context-induced changes in neural encoding, and situate computational mechanisms such as lateral inhibition and divisive gain control within a broader efficient coding framework^9^. Divisive normalization is considered a fast mechanism that operates within local populations of sensory neurons^4,7^. This is consistent with previous perceptual results showing that the tilt illusion follows dynamic changes of the surround orientation up to 10 Hz^47^. It is also consistent with our finding that surround modulation is spatially confined to ROIs covering the center-surround stimulus boundary. Previous behavioral studies of the tilt illusion further corroborate this by showing that stronger segmentation cues at the center-surround boundary decrease the strength of the illusion^48,49^.

Although our study was focused on characterizing the changes in sensory encoding, we demonstrate that these changes are sufficient to accurately predict subjects’ reports of their perceived tilt illusion using a recently proposed Bayesian observer model^18^. The specific model currently provides the most accurate quantitative descriptions of human behavior in orientation estimation tasks. Its predictions support the causal role of the encoding changes in creating the tilt illusion. It also suggests that the tilt illusion is not the result of sub-optimal inference processes but rather reflects resource-rational behavior in a statistically structured environment. It is also worth noting that the Bayesian observer model assumes that subjects’ reported orientation estimates are affected by an ordinal/categorical assessments of the stimulus, i.e., whether the orientation of the center stimulus is perceived to be clockwise or counter-clockwise of the surround orientation. This suggests that in addition to modulating encoding, the surround stimulus also acts as a reference in guiding subjects’ reports, which links the tilt illusion to contextual effects often referred to as reference repulsion (e.g., Treue et al.^50^). An implication is that the bias in reported orientation estimates seen is in part non-perceptual, arising from downstream decision processes. As a result, the repulsive biases in subjects’ reported estimates may exaggerate the actual perceptual distortions they experience with the tilt illusion.

Our results offer novel predictions regarding other aspects of contextual effects. In the case of the tilt illusion, previous research has shown that different surround features — such as complex textures with a broader range of orientations^51^ — can also induce the illusion, although with different magnitudes. We hypothesize that the perceptual characteristics of the illusion can be predicted based on the shape of the conditional orientation distribution. For instance, a surround containing a wider range of orientations is expected to predict a broader and more gradual increase in the probability of the center orientation, resulting in weaker change in encoding and, consequently, a smaller bias.

It is also worth considering the temporal homolog of the tilt illusion, i.e., the tilt aftereffect. In the tilt aftereffect, context is established temporally through a sequence of preceding stimuli with fixed orientation^52^. The changes in orientation perception and neural tuning observed for the tilt aftereffect are remarkably similar to those found in the tilt illusion^10,53,39^. Furthermore, the conditional orientation distribution for temporally adjacent stimulus is very similar to that of spatial contexts, also peaking at the dominant orientation of the context^10,39^. Therefore, we predict similar changes in encoding precision for the tilt aftereffect as we have reported here for the tilt illusion. We have recently shown that this is the case based on psychophysical threshold measurements^39^. It will be intriguing to further validate this using fMRI data and to investigate the extent to which the increase in contextual modulation along the visual ventral stream is also observed for the tilt aftereffect.

Finally, our methods provide a general framework for understanding context effects across other perceptual domains, including shape (e.g. the Ebbinghaus illusion^54^), motion^55^, color^56,57^, and face perception^58^. Our results suggest that these phenomena all originate from context dependent changes in sensory representation that reflect the context-conditioned statistics in natural visual environments.

## Acknowledgments

This work was funded by a Collaborative Research Grant from MindCORE of the University of Pennsylvania, P30EY001583, and R01EY036255 (to G.K.A.). L.-Q.Z. is supported by the Howard Hughes Medical Institute. J.M. is in part supported by NSF grant IIS-1912232 (to A.A.S.). We thank Huseyin O. Taskin and Marlie Tandoc for their assistance with fMRI data collection. We also thank Dr. Ruben van Bergen and Dr. Janneke Jehee for sharing their data. Some of the results described in this paper were presented at the Annual Meeting of the Vision Science Society in May 2024, and the Annual Conference on Cognitive Computational Neuroscience in August 2024.

## Methods

### Experiment

This study was approved by the University of Pennsylvania Institutional Review Board in accordance with the Declaration of Helsinki, and all participants provided a written consent.

### Procedure

Subjects (n = 10) performed a delayed orientation estimation task conducted in the fMRI scanner. All subjects had normal or corrected to normal visual acuity. On each trial, a 2 s initial delay was followed by the presentation of an oriented grating stimulus for 1.5 seconds. The oriented stimuli were presented within an annular surround of either non-oriented noise, or gratings with one of two fixed orientations (± 35 degrees off vertical). After a blank delay period of 4-5 s, a line probe appeared, and subjects used a two-button response pad to rotate the probe to report their orientation estimates. The line probe remained on the screen for a duration between 3.5 and 4.5 s long (uniformly sampled). The blank delay period was configured such that the total time of delay and response was 8.5 seconds. The visual stimulus and response task were created using PsychoPy^59^.

Each fMRI acquisition consisted of 20 trials, with all trials within the acquisition using either the non-oriented surround, or one of the two oriented surrounds. The assignment of surround condition to acquisition order was counterbalanced within and randomized across subjects. Over six sessions of fMRI scanning, subjects completed a total of 60 acquisitions, resulting in 1,200 trials (400 trials for each surround condition).

### Stimulus

Subjects viewed stimuli on an LCD monitor positioned at the end of the scanner bore via an angled mirror mounted on the head coil. Each stimulus consisted of a mid-gray central region with a radius of 1.5 deg, and a fixation dot of 0.35 degrees. An oriented grating target occupied the area between 1.5 and 7 degrees radius, and had a spatial frequency of 1 cycle per degree. The orientation was sampled uniformly between 0 and 180 degrees. Around the grating target was an annular surround extending from 7 to 12.5 deg radius. It contained either non-oriented noise, or one of the two fixed orientations (±35 deg), all with a spatial frequency matched to the center (1 cycle per degree). The entire stimulus was contrast-modulated at 1 Hz temporal frequency with a peak contrast of 20%. See Fig. 5A for a schematic of the spatial configuration of the stimulus.

### Neuroimaging

#### MRI acquisition

Anatomical (T1w and T2w) and Blood Oxygen Level Dependent (BOLD) functional images were acquired on a Siemens 3T Prisma scanner with a 64-channel head coil at the University of Pennsylvania. For T1w images, the tfl3d1 sequence was used with 0.8 mm isotropic voxels, TR = 2,400 ms, TE = 2.2 ms, and flip angle = 8 deg. For T2w images, the SPC sequence was used with 0.8 mm isotropic voxels, TR = 3,200 ms, TE = 563 ms, and flip angle = 120 deg. The functional images were acquired with the spin echo imaging sequence epfid2d1, with 2 mm isotropic voxel size, TR = 800 ms, TE = 37 ms, flip angle = 52 deg.

#### Retinotopic mapping

Each subject performed an additional scanning session devoted to retinotopic mapping. The stimulus consisted of a black and white checkerboard pattern that contrast-reversed at 5 Hz temporal frequency. This pattern was displayed against a mid-gray background within a circular aperture 21 degrees in diameter. The bar moved along both cardinal and oblique orientations, with the sequence of bar positions played in reverse for the second half of the acquisition. Subjects were instructed to focus on a central black fixation dot throughout the measurements and to respond with a button press when the dot occasionally and briefly turned red. Each acquisition was 330 sec, and each subject completed 6 acquisitions. T1w and T2w anatomical images were also acquired at the end of the retinotopic mapping session.

The retinotopic mapping data were analyzed using previously developed procedures^60^. Briefly, a noise removal method based on independent component analysis was first applied to the functional measurements^61,62^. Population receptive field (pRF) maps were then produced by fitting a model that jointly estimates the voxel pRF and hemodynamic response function^63,64^. Lastly, the pRF estimates were combined with the cortical surface topology derived from structural measurements within a Bayesian framework to produce a final retinotopic map for each subject^65^. The boundary of visual areas and the visual eccentricities of voxels were defined based on this map.

#### MRI data preprocessing

We processed both the structural and functional data using the Human Connectome Project (HCP) minimal processing pipeline^66^. This stage corrected for gradient nonlinearity, motion, and phase encoding direction in volumetric images. Subsequently, voxels were mapped onto a cortical surface template (fsaverage), with an additional 2 mm FWHM Gaussian surface smoothing applied. The resulting time series was high-pass filtered with a cutoff of 150 sec to remove slow drifts in the BOLD response, and linear regression against the motion regressors generated by the HCP pipeline was used to further remove motion artifacts. To obtain the voxel activity pattern for each stimulus presentation, the time series for each trial within a session was first aligned based on stimulus onset, normalized (z-score) across the corresponding time point, and averaged between 4 and 8 seconds.

#### Region of interest

We defined regions of interest (ROIs) based on the retinotopic maps obtained using the procedure described above. In our primary analysis, we selected voxels with pRF centers between 1 and 7 degrees of visual eccentricity, and from the following (groups of) visual areas: V1 + V2 + V3 (early visual cortex); V1 alone; V2 + V3; hV4 + VO1 + VO2; and V3A + V3B. Additionally, we established two control areas, the auditory cortex (A1 + A2) and primary motor cortex (M1) based on the cortical parcellation template produced by Glasser et al.^67^. In an alternative analysis, we expanded the voxel pRF center to the range of 1 to 15 degrees of visual eccentricity, covering the entire stimulus.

To understand the spatial profile of the surround modulation effect, we conducted an additional analysis in which voxels within area V1 - V3 were chosen based on their pRF center *c* and size *σ* in units of visual degrees (Fig. 5). To select voxels exclusively from within the center region, we defined two ROIs using the criteria *c* + 2*σ <* 1.5, and 1.5 *< c* + 2*σ <* 5. To select voxels exclusively from within the surround region, we defined two other ROIs with 9 *< c* − 2*σ <* 15, and 15 *< c* − 2*σ <* 30. Lastly, voxels at the center-surround boundary were selected as 5 *< c <* 9.

#### Theoretical framework

We modeled orientation perception as an encoding-decoding process^68^: Stimulus orientation *θ* is encoded as a noisy neural measurement *m*, described by the encoding model *p*(*m*|*θ*). Perceptual estimates 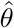 are then formed through a decoding process 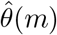 based on the neural measurement *m*. The Fisher Information (FI) of the encoding is defined as:

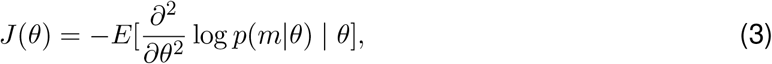

and quantifies the encoding accuracy as a function of *θ*. For a neural population that encodes information efficiently given limited encoding resources, there is a direct relationship between the stimulus prior distribution *p*(*θ*) and encoding accuracy *J*(*θ*)^16,17,69^:

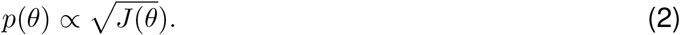

The goal of our analysis was to infer *J*(*θ*) independently from behavioral data (referred to as behavioral FI) and neural data (referred to as neural FI). We elaborate on the methods we used to derive these quantities in the sections below.

#### Behavioral data analysis

On each trial of the experiment, subjects produced an estimate 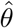 of the true stimulus orientation *θ*. For a given *θ* across trials, those estimates formed a distribution 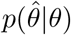. We denote the bias *b*(*θ*) and variance *σ*^2^(*θ*) of subjects’ estimates (Fig. 1) as

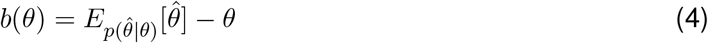

and

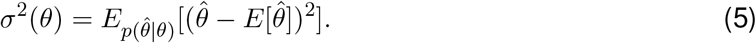

Both are defined as a function of *θ*. To compute these quantities from response data, we applied a sliding window analysis with a window size of 18 deg. The mean and variance were computed within each window with the true *θ* being the center of that window.

#### Cramer-Rao lower bound

Given an encoding model *p*(*m*|*θ*) with FI *J*(*θ*), the Cramer-Rao Lower Bound (CRLB) states that for an biased estimator 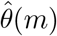 Fisher Information is bound from below^19^ as

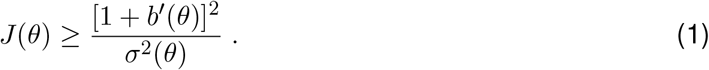

Here *b*^′^(*θ*) denotes the derivative of the bias *b*(*θ*). Thus, the Cramer-Rao bound specifies a lawful relationship between encoding accuracy and the bias and variance of an estimator^13,12^. To interpret Eq. 1, we can denote *g*(*θ*) as the mean estimate 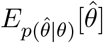. We have *b*(*θ*) = *g*(*θ*) − *θ*, and the inequality Eq. 1 can be expressed as

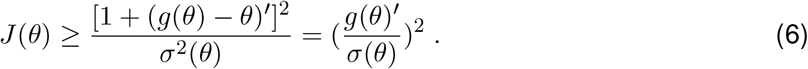

For an unbiased estimator *g*^′^(*θ*) = *θ*^′^ = 1. In this scenario, there is an inverse relationship between *J*(*θ*) and *σ*^2^(*θ*). When |*g*^′^(*θ*)| *<* 1, the estimator performs a local compression, leading to a reduction in variance. Conversely, if |*g*^′^(*θ*)| *>* 1, the estimator expands the local space, causing an increase in variance relative to 1*/J*(*θ*).

In our analysis, we assume the lower bound to be tight (or equally loose) for every *θ*. This allows us to infer FI from the measured estimation bias and variance. We have previously shown that a wide range of decoders, including those commonly used such as maximum likelihood and Bayesian decoders, all attain the lower bound^13^. We independently applied CRLB to the estimation data from the non-oriented and oriented surround conditions, obtaining two sets of behavioral FI curves for the baseline (Fig. 1D) and the surround modulation condition (Fig. 4B), respectively. The standard error (SEM) was estimated through a bootstrapping procedure that resampled the raw data 500 times.

Lastly, unless stated otherwise, we report the normalized, square root of FI throughout this article denoted as

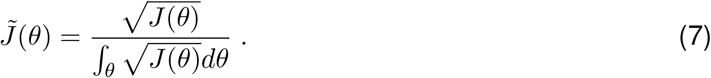

This facilitates the comparison of FI measured for different conditions, but also highlights the relationship between encoding precision and prior distribution as proposed by efficient coding (Eq. 2): 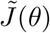 can be interpreted as the orientation prior for which the neural coding is most efficient. The denominator, 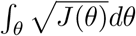, measures the amount of total encoding resources.

### Neural data analysis

#### Voxel encoding model

We modeled the voxel activity pattern **m** based on a probabilistic encoding model developed previously in Van Bergen et al.^14^. We denote this model as *p*(**m**|*θ*). The model starts by assuming a set of basis tuning functions in orientation space:

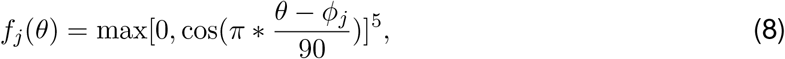

where *θ* is the stimulus orientation in degrees, *ϕ*_*j*_ denotes the orientation preference of the j-th function. We use *J* = 8 in our analysis, with the preferred orientation spaced equally between 0 and 180 degrees.

The activity of each voxel *m*_*i*_ was modeled as a weighted sum of the responses of the basis function:

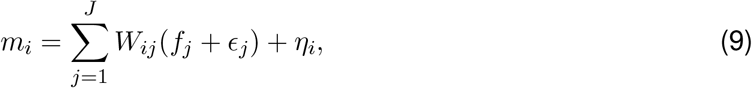

where *W* is the weight matrix. The model incorporates two sources of noise: each basis function is affected by independent channel noise *ϵ*_*j*_ with variance *σ*^2^: *ϵ*_*j*_ ∼ 𝒩 (0, *σ*^2^); and the residual noise in each voxel is modeled as *η* ∼ 𝒩 (0, Σ_*η*_). The residual covariance matrix is constructed as:

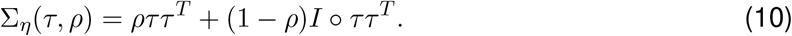

The diagonal terms of Σ_*η*_ are 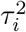, which represents the residual variance of each voxel *i*, whereas *ρ* is a global correlation parameter such that the off-diagonal terms of are *ρτ*_*i*_*τ*_*j*_.

Together, this model defines *p*(**m**|*θ*) as a multivariate normal distribution:

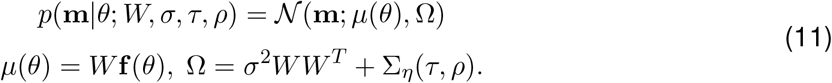

#### Model fitting

We fit separate encoding models to the voxel activity pattern obtained for every subject for each surround condition, and at each ROI. Each surround condition had 400 trials, with the number of voxels ranging from approximately 300 to under 2,000 depending on the ROI. A cross-validation procedure was employed in all cases, where the 400 trials were divided into 20 folds. One fold served as the hold-out data, while the model fitting was performed on the remaining folds. Orientation decoding and Fisher information estimation were only conducted on the held-out data. This process was iterated until each fold had become the hold-out data once. Lastly, to avoid potential biases introduced by the specific choice of basis function, four different encoding models with phase-shifted tuning curves were fit, and results were obtained by averaging across them.

The parameters of the encoding model were obtained using a two-step procedure^14^. The weight matrix was first estimated through ordinary linear regression. Denote matrix *X* ∈ ℛ^*N* ×*J*^ as the responses of *J* basis functions across *N* trials, and matrix *M* ∈ ℛ^*N* ×*K*^ as the activities of *K* voxels across *N* trials, we have:

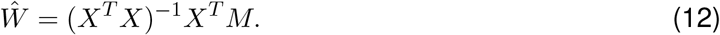

In the second step, the remaining noise parameters *σ, τ, ρ* were estimated using a maximum likelihood produce given a fixed *Ŵ* :

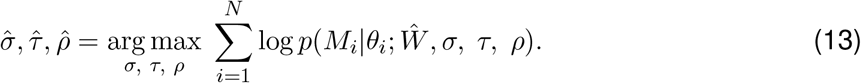

The encoding model was implemented in PyTorch^70^, and the maximum likelihood was performed using the sequential least squares programming algorithm in Scipy^71^. The model fittings are computationally expensive, but can be sped up significantly on GPUs with PyTorch.

#### Fisher information

For each trial in the held-out data with true stimulus orientation *θ*^*^ and voxel response **m**^*^, the orientation log-likelihood can be defined using the encoding model fitted to training data (Fig. 2B):

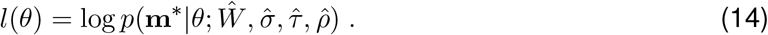

Orientation decoding was performed using the maximum likelihood decoder 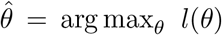 (see Fig. 2C - D). To obtain the neural Fisher information, we computed the negative second derivative of the log-likelihood function evaluated at *θ*^*^:

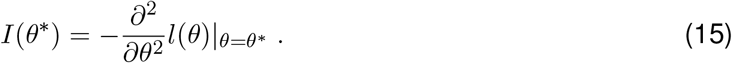

This quantity *I*(*θ*^*^) is called observed Fisher information^72^ (i.e., FI for a specific sample of **m**^*^), whereas the true Fisher information *J*(*θ*) is the expected value over *I*(*θ*): *J*(*θ*) = *E*_*m*_[*I*(*θ*)]. For each condition in our experiment, we obtained 400 estimates of observed FI *I*(*θ*) across orientations. The values for *J*(*θ*) and its standard error (SEM) were calculated by averaging *I*(*θ*) within a 25-degree window centered at various orientations (e.g., Fig. 2E).

Consistent with the behavioral data analysis, we report the normalized neural FI 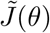 as defined in Eq. 7. The only exception was the calculation of the surround modulation index in Fig. 4D, Fig. 5C, and Fig. S3A. In these cases, we computed the difference between surround and baseline in the average, un-normalized *I*(*θ*) within a 25-degree window centered at the surround orientation (i.e., between 22.5 - 47.5 deg). This difference was then converted to a percentage change relative to the average *I*(*θ*) across all orientations in the baseline. Statistical significance was assessed using an unpaired t-test on the *I*(*θ*) samples within this 25-degree window.

#### Observer model for orientation estimation

We predicted bias and standard deviation of subjects’ perceived orientation reports using a recently proposed Bayesian observer model^18^. In the following we provide a compressed description of the model, and refer the reader to the original article for additional details.

The model assumes that orientation encoding is efficient based on the statistical (prior) distribution *p*(*θ*) over orientation *θ* in the observer’s environment (Eq. 2). Moreover, it assumes that perception and the downstream decision and control process operate *holistically* on all levels of the representational hierarchy; here this includes a higher, categorical representation of orientation *C* (e.g., cardinal vs. oblique orientations) in addition to the feature level representation *θ*. Thus, the model assumes that based on a sensory signal *m* the observer infers posterior beliefs at both levels of the hierarchy, i.e., *p*(*θ*|*m*) and *p*(*C*|*m*), which then provide the information for the downstream decision processes.

The orientation estimation task of our experiment requires the observer to adjust a probe stimulus such that its orientation matches the perceived orientation of the center grating (test) (Fig. 1A). The model assumes that the observer infers the posterior beliefs of both the orientation and the category for each of the two stimuli, probe and test. As the observer adjusts the orientation of the probe, they seek to report the probe orientation *θ*_*p*_ that minimizes the expectation of a joint objective *L*_tot_ that reflects the mismatch between the two stimuli at both the feature and the category level; hence

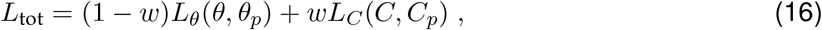

where *L*_*θ*_ is defined as the cosine difference between the test and the probe orientation, and *L*_*C*_ is a fixed cost if test and the probe stimuli fall into different orientation categories but zero otherwise.

For the model simulations (Fig. 6, encoding precision and the orientation prior used for Bayesian inference were determined by the neural FI of areas hV4/VO1/VO2 (Fig. 4) measured in the baseline (non-oriented surround) and the oriented surround condition, respectively. In the baseline condition, we closely followed the model specifications of the original study, assuming two orientation categories (clockwise and counterclockwise relative to vertical), and parameter values for category overlap *κ*_*card*_ and boundary noise *κ*_*b*_ similar to the values in Mao and Stocker^18^. The encoding noise *κ*_*i*_, the weight *w* of the categorical mismatch, and an additive motor noise *κ*_*m*_ were adjusted so that the magnitude of the bias and standard deviation matched the data in the baseline condition. We then predicted behavior in the oriented surround condition (tilt illusion) based on this model, further assuming that the surround orientation created an additional category boundary with relative sharp boundaries (high *κ*_*surr*_) as the surround is always present. The following table list the values of all model parameters for simulating the tilt illusion:

**Table 1:** Model parameters used for the simulations shown in Fig. 6.

| Parameter | Value |
| --- | --- |
| $\kappa_i$: sensory noise | 10.5 |
| $\kappa_b$: boundary noise | 60 |
| $\kappa_{card}$: cardinal category overlap | 2 |
| $\kappa_{sur}$: surround category overlap | 24 |
| $w$: categorical weight | 0.72 |
| $\kappa_m$: motor noise | 48 |

#### Voxel encoding model based on steerable pyramid

We estimated the changes in the measured encoding FI that arise only due to differences in stimulus configuration (i.e., non-oriented vs. oriented surround) in the absence of any potential change in neural responses properties. We follow the approach of Roth et al.^33^ to create an image-computable model of voxel encoding (Fig. 7A). For a given stimulus image we use the steerable pyramid^34^ to create filtered responses at different orientations (*c*) and spatial frequency (SF) bands (*s*). We used a complex pyramid and combined the real and imaginary parts to obtain single energy-like filter responses. This yielded multiple filtered images indexed by *c* and *s*: *r*(*c, s*). These images can be thought of as representing V1-like neuronal responses at every location of the visual space, each with different orientation and spatial frequency selectivity.

To simulate voxel activity *f* (*θ*), we combined responses across these orientation and SF bands as *f* (*θ*) = Σ_*c,s*_ *r*(*c, s*). This produced a final “retinotopic map” of voxel responses. In general, each band *r* can be weighted differently, resulting in voxel selectivity over orientation and spatial frequency. Here, we used equal weights as we are only interested in the difference between two stimulus conditions. The encoding FI is defined as 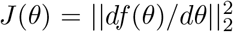, which is the FI assuming independent Gaussian response noise with unit variance for each voxel.

In our case, a steerable pyramid with 6 orientation and 6 SF bands was constructed for each stimulus image (using Pyrtools^73^). The final response map *f* (*θ*) was obtained by averaging over all orientation bands and SF bands 3, 4, and 5, as these SF bands exhibit the largest (worst-case) changes in FI. To combine different SF bands, we downsampled response maps at finer scales to match the resolution of the coarser scale. To compute the changes in FI, we first computed *J*(*θ*)_base_ using stimulus images with non-oriented surround. Note that *J*(*θ*)_base_ is non-zero, representing the vignetting effect reported by Roth et al.^33^, and is also uniform, since any effect of stimulus configuration in the non-oriented condition is isotropic by construction. We then computed *J*(*θ*)_surr_ using stimuli with oriented surround. The change in was is calculated as Δ*J*(*θ*) = *J*(*θ*)_surr_ − *J*(*θ*)_base_, and the results are shown in Fig. 7B. See Supplementary Fig. S5 and the associated text for a more extensive discussion on the issue of stimulus vignetting, including FI calculated separately at each spatial scale.

## Code and data availability

The behavioral data and the preprocessed fMRI data from this study can be accessed through the Open Science Framework: https://osf.io/9uqbd. The raw fMRI data are available upon request. The software code developed for data analysis is available through GitHub: https://github.com/lingqiz/orientation-encoding.

## Supplementary Information

### Supplementary figures

**Supplementary Figure 1:**
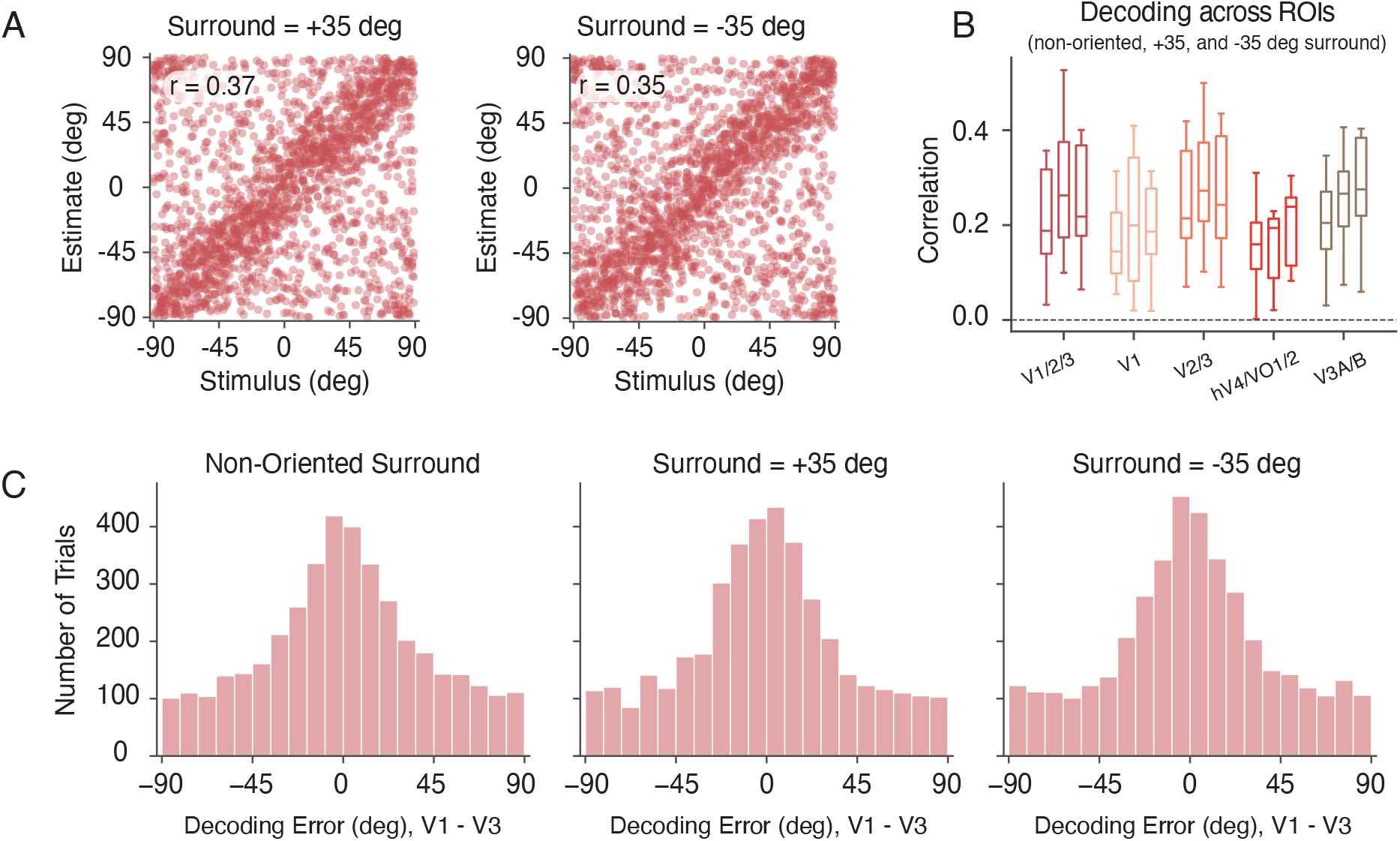
Orientation decoding performance. **A)** Same as Fig. 2C, scatter plot of the stimuli orientation (x-axis) versus the decoded orientation (y-axis) from the early visual cortex (V1 to V3), for the two oriented surround (± 35) conditions. **B)** Same as Fig. 2D, but with decoding correlation plotted separately for each of the three surround conditions within each ROI. **C)** Histogram of decoding errors from V1 - V3, for the combined subject across the three surround conditions.

**Supplementary Figure 2:**
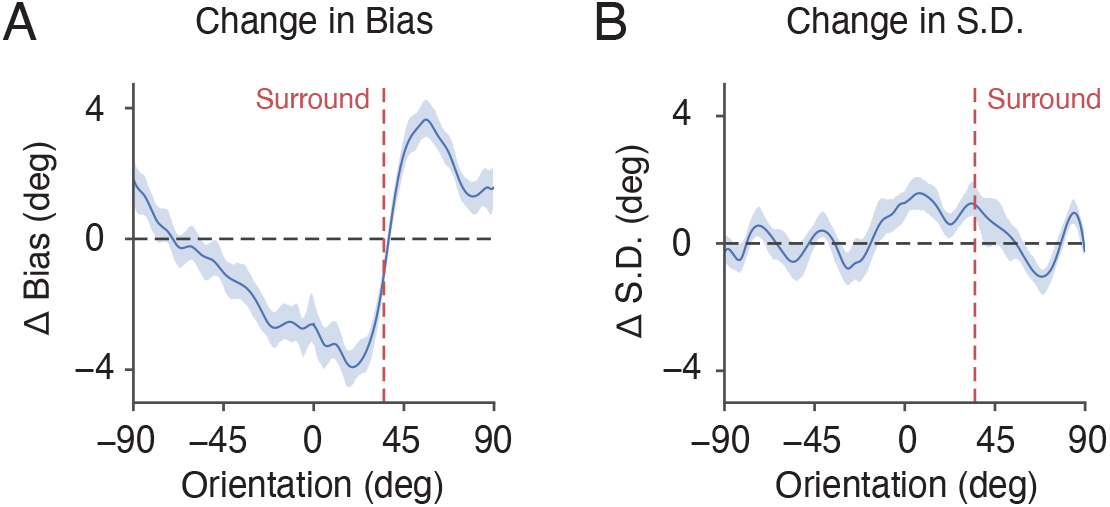
Effect of surround modulation on orientation estimation. **A)** Difference in estimation bias between the non-oriented surround and the oriented surround condition. **B)** Difference in the standard deviation of the orientation estimates between the non-oriented surround and the oriented surround condition. The shaded area indicates ±SEM.

**Supplementary Figure 3:**
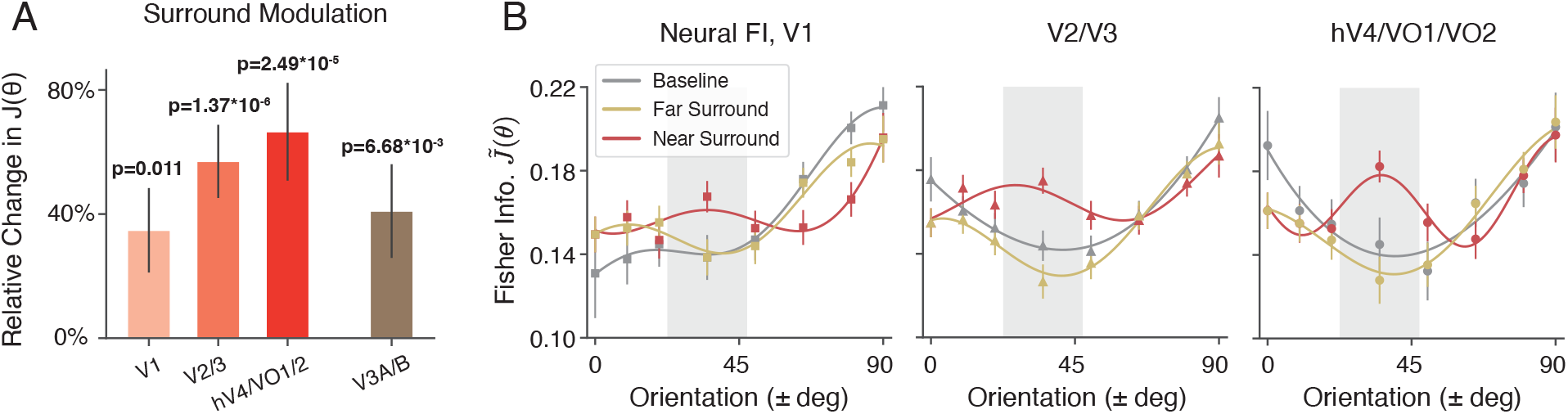
Neural encoding across visual areas with expanded eccentricity ROI. We repeated the same analysis as in Fig. 4D - E, but expanded the eccentricity selection to between 1 and 15 degrees to cover the entire stimulus. **A)** The relative change in neural FI with respect to the baseline near the surround orientation across different visual cortex ROIs. **B)** Comparison of neural FI along the visual ventral stream, between the near-surround side, far-surround side, and the baseline condition. Error bars indicate ±SEM.

**Supplementary Figure 4:**
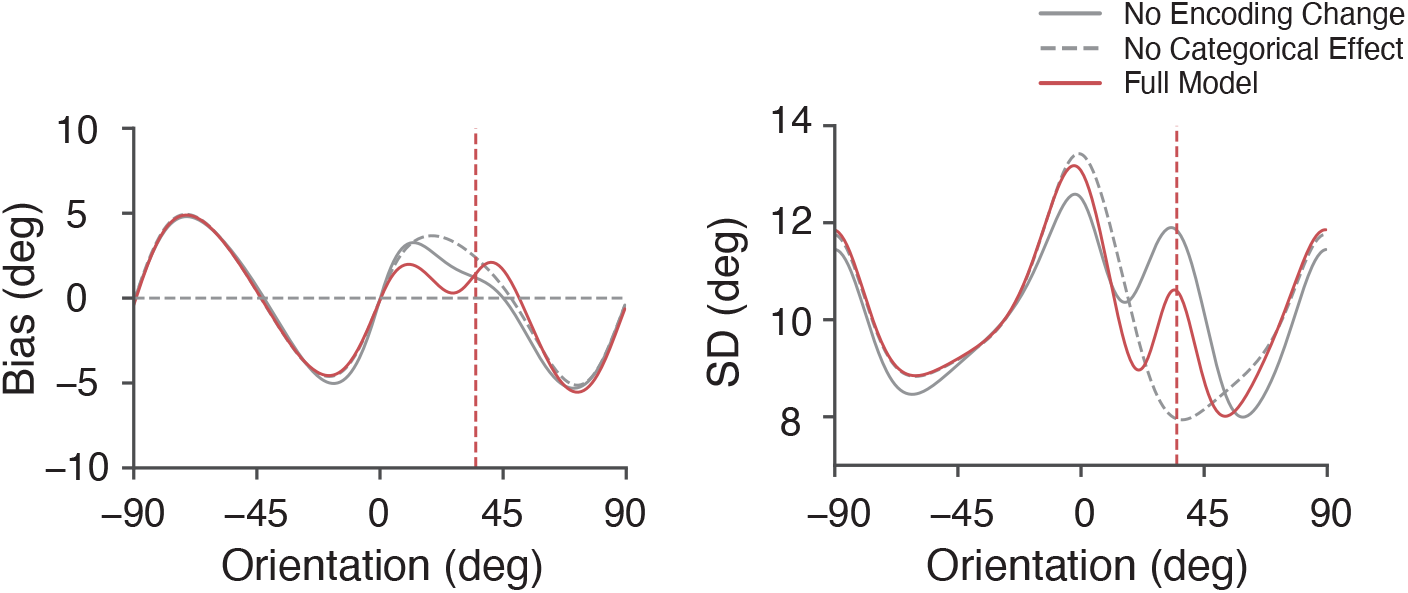
Both the dynamic change in sensory encoding and the categorical boundary are necessary for the correct model prediction of the tilt illusion. Panels show the predicted estimation bias and standard deviation of the observer model in the surround condition: Solid gray lines represent the model prediction without assuming a change in sensory encoding (i.e. using the encoding pattern from the baseline condition), while the dashed gray lines represent the model prediction without assuming the categorical boundary at the surround orientation. The solid red lines represent the prediction based on the full model (same as in Fig. 6). Both mechanism are required to correctly predicts the characteristic repulsive bias in the tilt illusion.

**Supplementary Figure 5:**
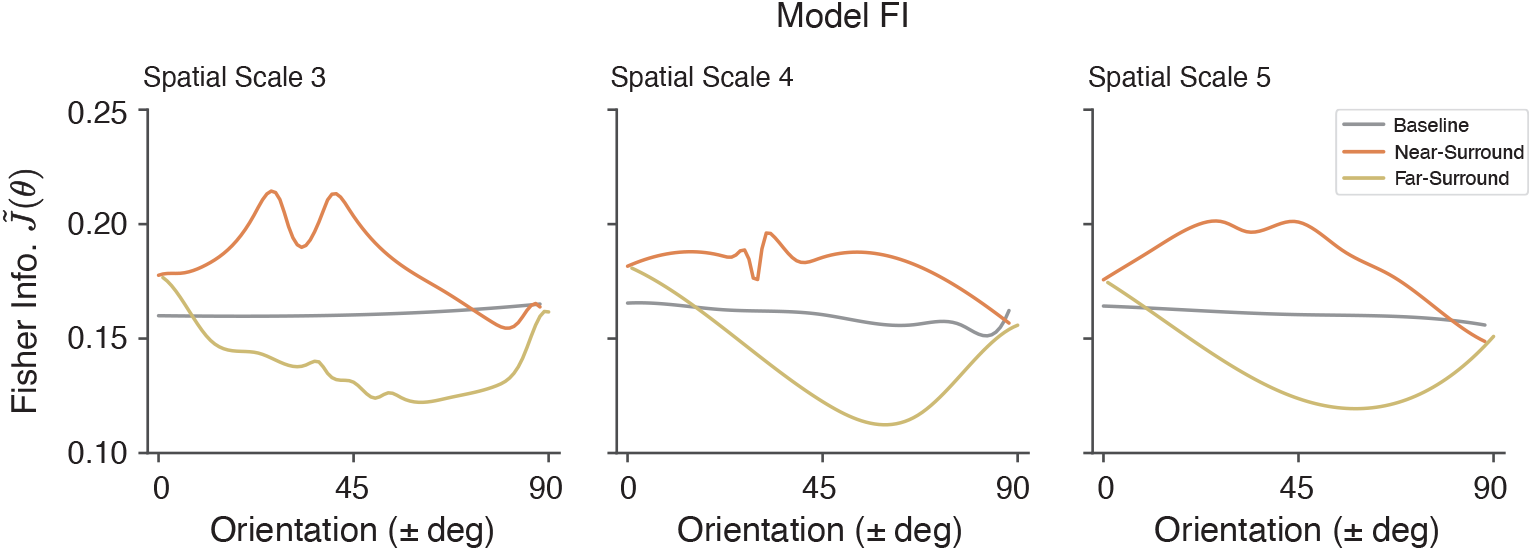
The effect of stimulus configuration on encoding FI. The neural basis of orientation decoding using functional imaging has been the subject of ongoing debate^32,33,74^, with recent findings challenging the notion that decoding is based on sensitivity to columnar-level neural tuning. In one sense, our results are independent of the outcome of this debate: Our neural measures of orientation encoding show strong consistency with behavioral data, indicating that regardless of the precise source of the orientation signal, it is indeed utilized by downstream processes and reflected in behavior. Furthermore, there are several notable features in our data that cannot be fully explained by stimulus and aperture configuration (i.e., vignetting) alone. In this analysis, we calculated the encoding FI of the voxel encoding model based on steerable pyramid decomposition at different spatial scales. The model is identical to that shown in Fig. 7A, except the voxel responses are averaged over orientation channels at a single scale. The encoding FIs are qualitatively similar in all cases: flat for the baseline, with a broad increase for near-surround orientations, and a broad decrease for far-surround orientations. There are at least three aspects of our data that are inconsistent with this “vignetting only” model. First, we observed an anisotropy in orientation encoding under the non-oriented surround condition. Given that the stimuli were designed to be isotropic (gray line), this effect must arise from anisotropies inherent in the neural representation of orientation. Second, we found that the effects of stimulus configuration in the oriented surround condition are broad and symmetrical at the surround and orientation orthogonal to it (orange and yellow line), inconsistent with the local changes we observed. Third, the model fails to replicate the increased effects of surround modulation across the visual hierarchy, as the effects of stimulus configuration remain similar across spatial scales. Therefore, while we do not rule out the possibility that stimulus configuration partially contributes to the tilt illusion, additional mechanisms in neural coding are necessary to fully explain our results.

### Supplementary data for individual subject

**Supplementary Figure 6:**
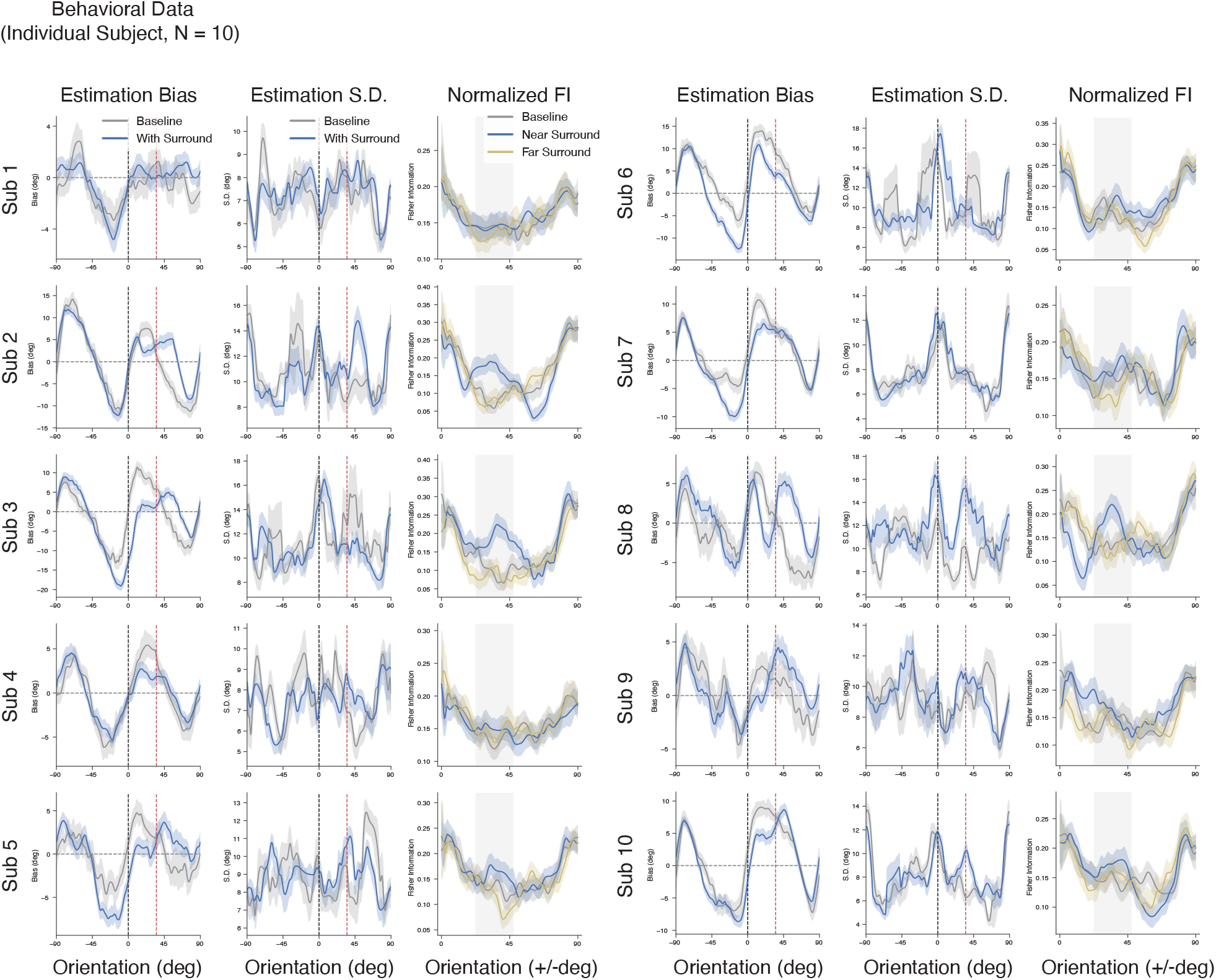
Behavioral data for individual subject. The bias, standard deviation of the orientation estimates, and the normalized behavioral FI for individual subject (N=10).

**Supplementary Figure 7:**
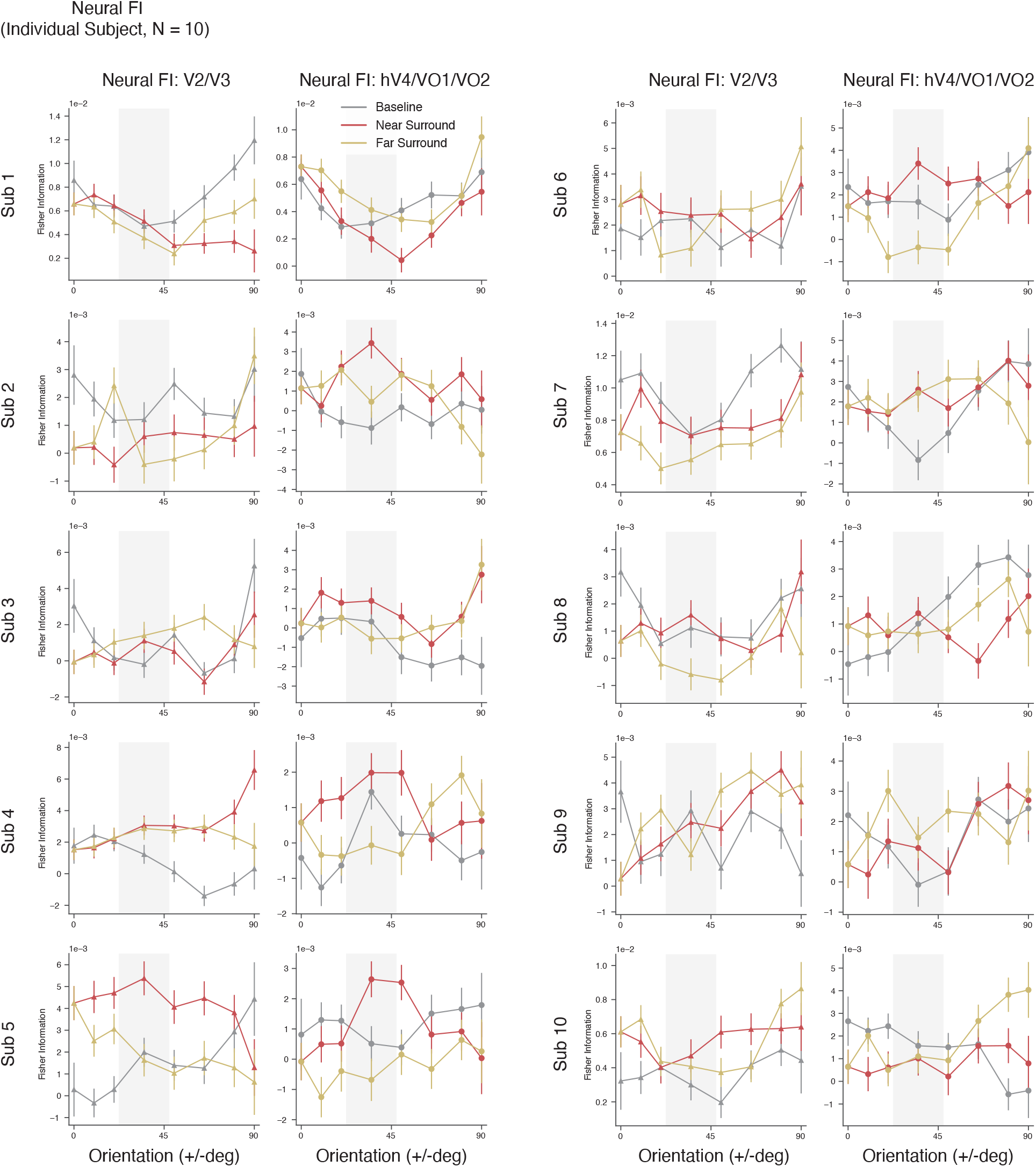
Neural FI for individual subject. The unnormalized neural FI from two visual area ROIs (between 1 - 7 degrees for V2/V3 and hV4/VO1/2) for individual subject (N=10).

